# The Role of Syllabic Rhythm in Speech Perception Across Languages

**DOI:** 10.1101/2025.03.18.643971

**Authors:** Irene de la Cruz-Pavía, Julián Villegas, Caroline Nallet, Kyoji Iwamoto, Ramón Guevara, Judit Gervain

## Abstract

The insertion of silences at regular intervals restores the intelligibility of English utterances that have been accelerated beyond comprehension, as long as the duration of the resulting speech-silence chunks falls within the theta rhythm of natural speech, i.e. the temporal modulation associated to the syllabic rate. We test whether such a rhythmic strategy works in languages rhythmically different from English, a stress-timed language. Thus, we assess whether comprehension of time-compressed Semantically Unpredictable Sentences (SUS) is restored in the syllable-timed language French and the mora-timed language Japanese, when silences re-establishing theta rhythm are inserted. Restoring the theta rhythm also improved intelligibility in French, but not in Japanese, in which best performance was instead achieved at faster rhythms, which suggests that modulation at the rate of a language’s basic rhythmic unit plays a key role in understanding speech. In a second experiment, French speakers listened to SUS with speech-silence chunks adapted to the range of the temporal modulations of the delta, gamma, and high gamma rhythms, which correspond to the rate of prosodic phrases, phonemes, and subsegmental features, respectively. Unlike the theta rhythm, we found no restorative effects, providing further evidence for the special status of the theta rhythm in speech comprehension.

## Introduction

A highly influential current theory of speech perception^1–3^ proposes that a hierarchy of nested neural oscillations in the auditory cortex operates at frequency bands that match the frequencies of certain linguistic units, explaining how the human brain processes speech at different levels from phonemes and syllables to words and phrases simultaneously. Specifically, delta oscillations (< 3 Hz) match the frequencies of prosodic phrases, theta oscillations (4–8 Hz) match the rate of syllables, while gamma oscillations (>30 Hz) match the rate of phonemes and subphonemic units^3–5^. These oscillations are aligned through a nesting relationship, whereby the phase of slower oscillations modulates the amplitude of faster oscillations^6^, thus ensuring that the different linguistic units are processed synchronously.

Modulations at the syllabic rate are particularly prominent in the speech signal. Indeed, speech in a large number of different languages shows the strongest amplitude modulation at a rate of about 4-5Hz^7,8^, i.e. within the theta range. These slow modulations are sufficient for adult and even young infant listeners to comprehend speech in silence^9,10^ and, according to the Rhythmic Segmentation Hypothesis proposed by Cutler^11^, they serve as the basic units for segmenting speech.

As a neural underpinning of this ability, the auditory cortex entrains to the speech envelope, i.e. the slow amplitude modulations of the speech signal, from birth^12–14^, showing high oscillatory power in the theta frequency band, comprised between 4–8 Hz^15,16^.

In striking illustration of the importance of the syllabic rate and the related theta rhythm, Ghitza and Greenberg^17^ elegantly demonstrated that accelerated speech compressed to a degree that adult listeners no longer comprehend becomes intelligible again if silence periods are systematically inserted to restore the theta rhythm. The authors presented participants with speech in English compressed to a third of its original duration, an acceleration that is no longer intelligible. They then chunked the 3x accelerated speech into segments of 40 ms, and added silences in between the chunks periodically or aperiodically. For the periodic stimuli, 6 conditions were created, which differed in the duration of the silence intervals: 0 ms (the original accelerated stimuli), 20 ms, 40 ms, 80 ms, 120 ms, and 160 ms. For aperiodic stimuli, the duration of the silence intervals varied pseudo-randomly around a mean corresponding to each of the periodic duration conditions. Participants’ comprehension accuracy of semantically unpredictable sentences (SUS; grammatically correct sentences without meaning, e.g. *The trip talked in the old stage*, changed in a U-shaped manner, being relatively impaired at the shortest and longest silence intervals (∼50% error), but improving when silences of medium duration (20 to 120 ms) were inserted, with the best comprehension results (∼20% error) for the periodic insertion of silences of 80 ms. Silences provide no additional linguistic or acoustic information, so it is not clear why they may aid comprehension at all. If a memory- or processing-time-based account is considered, e.g. if silences provide additional time for processing or for recall, then comprehension accuracy should increase linearly with silence duration. However, a U-shaped performance was found. The authors argue that this may be best explained by the endogenous rhythms of the auditory cortex, as the addition of silences between 20–120 ms restores a rhythm that is close to the theta rhythm of natural speech. Indeed, adding 80 ms of silence to a 40 ms accelerated chunk restores the original 120 ms duration of the uncompressed speech segment, such that speech rate becomes optimally fitted again to the syllabic rate of the original.

This is an intriguing demonstration of the importance of the syllabic rate in speech processing. However, it has only been tested in English so far. Yet, the languages of the world differ in their rhythms both at the linguistic level^18–22^ as well as at the level of temporal modulations in the speech signal^7^. Is the syllable rate equally relevant for the processing of different languages? Or would different languages have different privileged units of processing?

Synergistically combining the embedded neural oscillations model^3^ and the Rhythmic Segmentation Hypothesis^11^, the current study sets out to answer this question by testing whether a similar U-shaped restoration pattern is found when adult native listeners comprehend accelerated speech in French and Japanese, in a paradigm similar to the one used originally by Ghitza and Greenberg^17^ with English natives.

English, French and Japanese differ in their rhythmic properties. English is a stress-timed language, French is syllable-timed, while Japanese is mora-timed. This classification is based on the relative proportions of vowels and consonants in the speech signal (%V) as well as the variability in the duration of consonant clusters^20^ (ΔC; although other metrics exist to characterize speech rhythm^18,19,21,22^). Japanese has a relatively large proportion of vowels (53%) in its speech signal, and consonant clusters are infrequent. English has a much lower proportion of vowels (40%) and highly variable consonant clusters. French is between Japanese and English for both measures^20^. Related to these rhythm differences, the amplitude modulation rate of the speech signals in the three languages are different^7^. The amplitude of the signal is modulated at ∼5.1 Hz in French, ∼4.9 Hz in Japanese and at ∼4.3 Hz in English.

The present study has three goals. First, we sought to determine whether restoring the syllabic rate (theta rhythm) of accelerated speech via pause insertion led to a similar increase in intelligibility in rhythmically different languages. To that end, in Experiment 1 we presented French and Japanese natives with a paradigm similar to the one in Ghitza and Greenberg^17^. We only tested periodic silences, as Ghitza and Greeberg’s^17^ results indicated better overall performance with periodic than with aperiodic silence rates. We hypothesized that restoring the syllabic rhythm within the theta range may lead to improved performance in French, given its syllable-timed rhythm and the wealth of experimental evidence suggesting that the syllable is the preferred unit of speech perception in this language in infants^23,24^ and in adults^25,26^. Less evidence is available about Japanese, but existing studies suggest that in accordance with this language’s mora-timed rhythm, the basic unit of speech segmentation in this language is the mora, a subsyllabic unit, and not the syllable^27^. We thus hypothesized that restoring theta rhythm may not be useful for Japanese participants, and best performance may be achieved at somewhat faster rhythms, i.e. by the insertion of shorter silences, as the mora is shorter than the syllable. We thus predicted that highest performance will occur for each language at the silence durations that best restore the language’s basic rhythm, in accordance with the Rhythmic Segmentation Hypothesis^11^. Note that we carry over the term “theta rhythm” from the neural oscillation literature as a metaphor of the optimal range for syllabic processing, as the present behavioural study does not measure neural activity.

Second, in Experiment 2 we examined whether and if yes, how the restoration of other speech rhythms^1,3,28^, e.g. the slower delta rhythm (1–3 Hz) corresponding to larger prosodic phrases or the faster gamma rhythm (>30 Hz) tracking (sub)phonemic units, may impact comprehension. Acoustically, these temporal modulations are less prominent in the speech signal than the syllabic rate^7,8^, but they have clear linguistic relevance. If the rhythmic segmentation hypothesis is true, then we do not expect strong gains when these frequencies are restored. Such a result would also help exclude the interpretation that the U-shaped pattern found by Ghitza and Greenberg^17^ was some kind of general “Goldilocks effect”, whereby the best performance is observed for non-extreme values. If the U-shaped response is found only for the theta rhythm, but not for delta or gamma, that provides evidence for the specificity of the effect.

Third, we also sought to extend Ghitza and Greenberg’s^17^ original findings to a larger group of participants, as their original sample consisted of only 5 listeners.

We, therefore, presented one group of Japanese and four groups of French speakers (n = 24 per group) with 48 SUS (e.g. Japanese: *Sonna e ga hen ni kogu* “That picture rows strangely”; French: *La salle marque l’or qui vole* “The room marks the gold that flies”). The SUS had been time-compressed by a factor of 3, cut into chunks of constant length that varied as a function of the group, and had silences inserted after each chunk (see Figure 1 and Table 1). Chunk duration was chosen so that its original, uncompressed duration fell within one of the following oscillatory bands: 160 ms in delta, 40 ms in theta, 8 ms in gamma, and 4 ms in high gamma. For instance, 3x compressed chunks in delta had a duration of 160 ms. The duration of its original, uncompressed version was thus 480 ms or 2.08 Hz, falling within the range of the Delta oscillatory band (1-3 Hz). For French, we tested each of these chunk durations for a different group of speakers, resulting in 4 groups. For Japanese, only the critical theta duration was tested, i.e. one group of participants. Following Ghitza and Greenberg^17^, we generated five periodic silence duration conditions per group, whereby the inserted silence periods had 0.5, 1, 2, 3, and 4 times the duration of the chunk duration, in addition to a baseline condition with no silences added.

**Figure 1.**
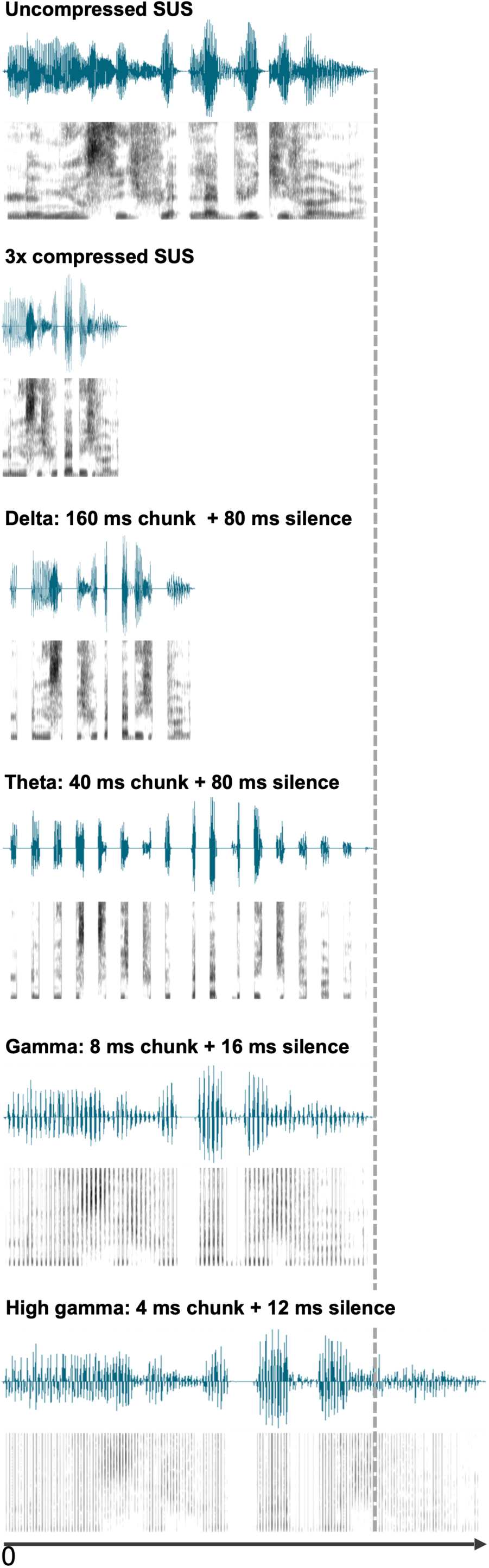
The SUS *Le mur siffle la buée qui vole* (“The wall hisses the mist that flies”) in six versions (top to bottom): (1) the original, uncompressed SUS, (2) time compressed by a factor of three, (3) split into 160 ms chunks with interleaved 80 ms silences, resulting in half the duration of the original uncompressed SUS (condition 2 of the delta group), (4) split into 40 ms chunks with interleaved 80 ms silences, resulting in the same duration as the original SUS (condition 4 of the theta group), (5) split into 8 ms chunks with interleaved 16 ms silences, resulting in the same duration as the original SUS (condition 4 of the gamma group), and (6) split into 4 ms chunks with interleaved 12 ms silences, resulting in 1.3 times the duration of the original SUS (condition 5 of the high gamma group). Each panel depicts the waveform (top) and spectrogram (bottom) of the SUS.

**Table 1.**
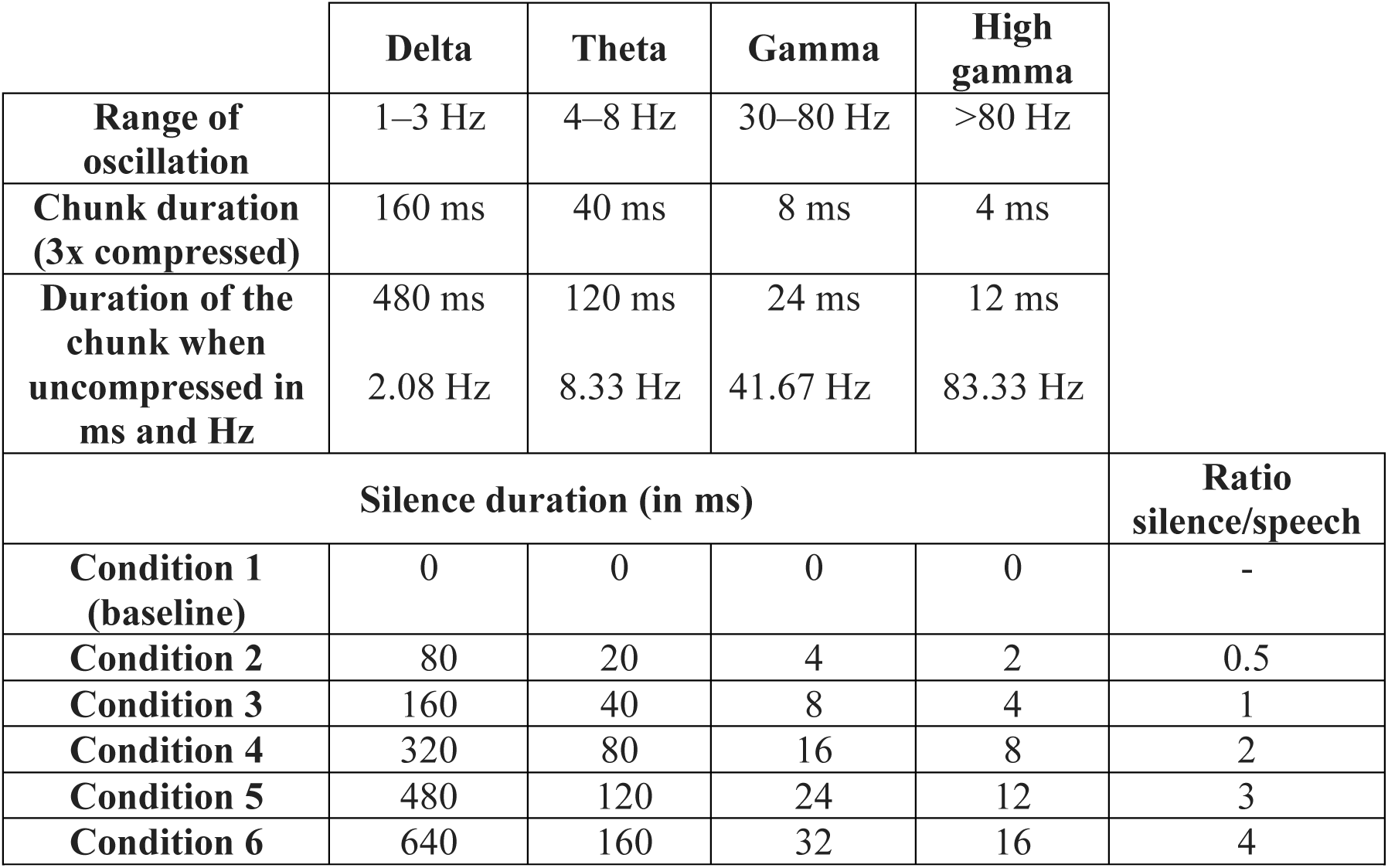
The table lists (from top to bottom): (a) the frequency ranges characteristic of the delta, theta, gamma and high gamma oscillations, (b) the duration of the compressed speech chunks in each of the groups tested, (c) the equivalence in Hz of the compressed chunks when uncompressed, showing that the uncompressed duration of the chunks in (b) fell within its corresponding oscillatory band, (d) silence duration in each of the 6 conditions tested per group, and finally the relative proportion of silence with respect to its corresponding speech chunk (rightmost column).

|  | **Delta** | **Theta** | **Gamma** | **High gamma** |  |
| --- | --- | --- | --- | --- | --- |
| **Range of oscillation** | 1–3 Hz | 4–8 Hz | 30–80 Hz | >80 Hz |  |
| **Chunk duration (3x compressed)** | 160 ms | 40 ms | 8 ms | 4 ms |  |
| **Duration of the chunk when uncompressed in ms and Hz** | 480 ms<br>2.08 Hz | 120 ms<br>8.33 Hz | 24 ms<br>41.67 Hz | 12 ms<br>83.33 Hz |  |
| **Silence duration (in ms)** |  |  |  |  | **Ratio silence/speech** |
| **Condition 1 (baseline)** | 0 | 0 | 0 | 0 | - |
| **Condition 2** | 80 | 20 | 4 | 2 | 0.5 |
| **Condition 3** | 160 | 40 | 8 | 4 | 1 |
| **Condition 4** | 320 | 80 | 16 | 8 | 2 |
| **Condition 5** | 480 | 120 | 24 | 12 | 3 |
| **Condition 6** | 640 | 160 | 32 | 16 | 4 |

After a brief training on the task, participants listened to 100 semantically meaningful sentences that were 3x compressed so that they had time to perceptually adjust to compressed speech. This has been shown to be necessary even for compression rates that are intelligible^29–31^. Immediately after, they listened to the 48 SUS (8 per each silence duration condition, each SUS appeared only in a single condition per participant), and were instructed to type whatever they had understood after each SUS. We calculated the percentage of correctly identified syllables per SUS (as well as morae in the Japanese group).

## Results

### Experiment 1. Restoring the theta rhythm in French vs Japanese

We analysed the responses of a group of French and a group of Japanese speakers (n = 24 each) presented with time-compressed and chunked SUS with the same chunk and silence durations as in Ghitza and Greenberg^17^. The percentage of correctly identified syllables are shown for each condition in Figure 2 and Table S1 of the supplementary material. Note that we report percentage of correct syllables, whereas Ghitza and Greenberg^17^ reported percentage of errors. Therefore, Ghitza and Greenberg’s^17^ results map onto an *inverted* U-shaped pattern in our presentation of the data.

**Figure 2.**
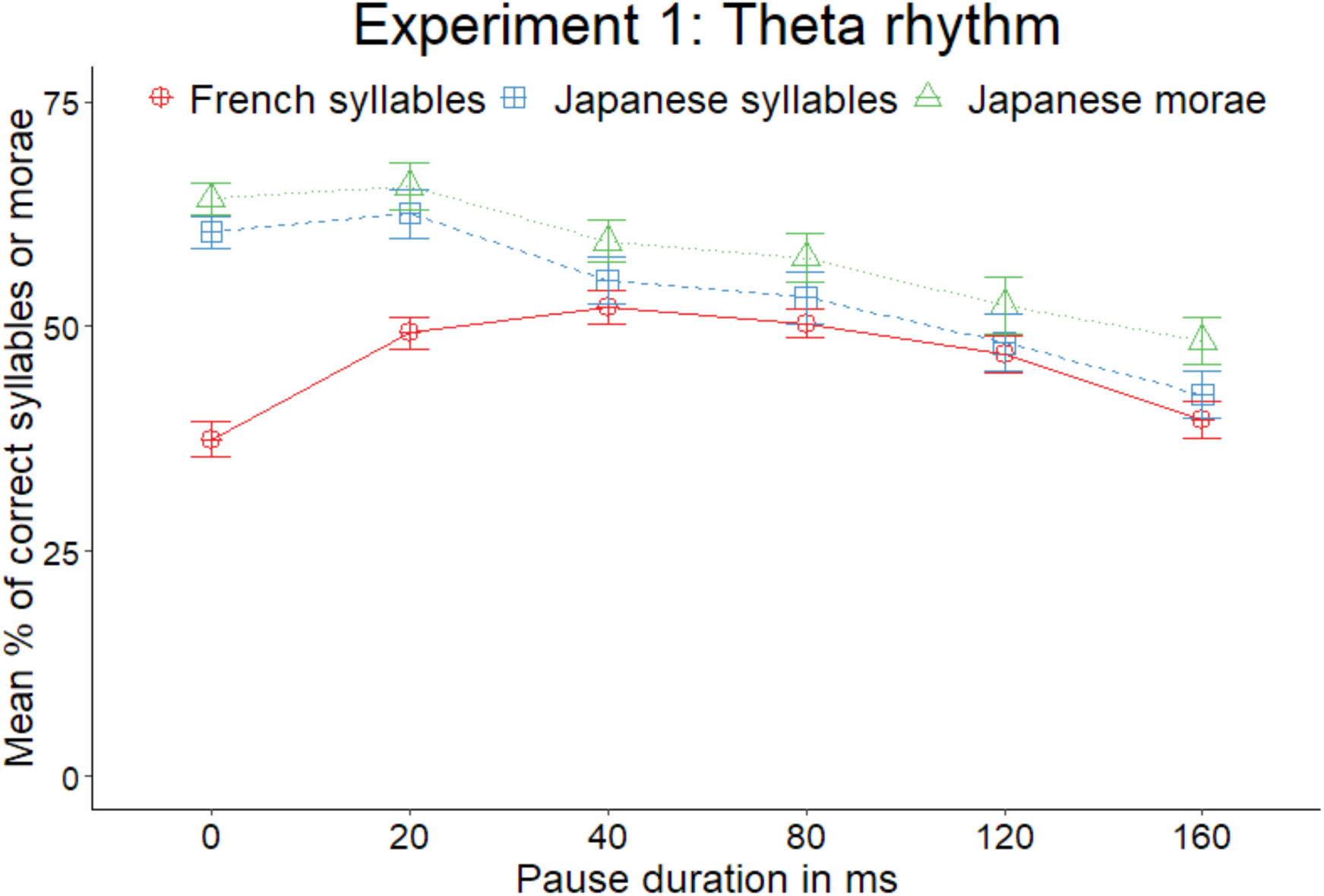
Accuracy by group and pause length. Each group’s mean accuracy is presented as a separate line. The y-axis depicts the mean percentage of correct syllables per SUS. The x-axis presents the 6 silence length conditions depicting their absolute temporal durations (in ms).

A mixed ANOVA with Group (French theta, Japanese theta) as a between-subjects factor and Pause Length (Conditions 1 to 6) as a within-subjects factor yielded a significant main effect of Group (*F*(1,46) = 11.0, *p* < .001, *η_p_²* = .107), of Pause Length (*F*(5,230) = 17.4, *p* < .001, *η_p_²* = .158), and a significant interaction between the two factors (*F*(5,230) = 11.8, *p* < .001, *η_p_²* = .113). Holm corrected pairwise post hoc tests revealed that accuracy in conditions 1 (i.e. the baseline) and 2 was significantly higher in Japanese as compared with French (both *p* < .001). The remaining comparisons did not reach significance (all *p* ≥ .353). Given the main effect of Group, due to greater overall accuracy in Japanese than in French, we ran one-way repeated-measures ANOVAs examining the role of Pause Length within each group. Both groups yielded a significant main effect of Pause Length (French theta: *F*(5,115) = 12.8, *p* < .001, *η²* = .268; Japanese theta: *F*(5,115) = 16, *p* < .001, *η²* = .225). Holm corrected pairwise post hoc tests yielded the following results. The analysis of the French theta group revealed that accuracy was significantly lower in the baseline than in all other conditions (all *p* ≤ .006) except for condition 6. Accuracy was also significantly greater in conditions 2, 3, and 4 than in condition 6 (all *p* ≤ .005). These results thus pattern into an inverted U-shape. Post-hoc analyses of the Japanese theta group revealed that accuracy was significantly better in conditions 1–4 than in condition 6 (all *p* ≤ .048). Accuracy was also significantly better in conditions 1–2 than in condition 5 (both *p* ≤ .015).

As morae and not syllables are argued to be the minimal unit of perception in Japanese^27^, we ran the same analysis coding accuracy as percentage of correct morae, which revealed a very similar pattern of results. A one-way repeated-measures ANOVA yielded a significant effect of Pause Length (*F*(5,115) = 12.9, *p* < .001, *η²* = .194). Post-hoc tests with Holm corrections revealed that accuracy was significantly better in conditions 1–3 than in condition 6 (all *p* ≤ .033). Accuracy was also significantly better in conditions 1–2 than in condition 5 (*p* ≤ .018).

In sum, the Japanese theta group displayed a pattern of decreasing intelligibility as silence duration increased regardless of the unit of measurement (syllable or mora), in contrast with the inverted U-shape obtained in the French theta group.

### Experiment 2. Restoring the delta, gamma, and high gamma rhythms in French

We analysed the responses of three groups of French participants (n = 24 each), presented with the delta, gamma and high gamma sets of silence conditions. The percentage of correctly identified syllables are shown for each condition in Figure 3 and Table S1 in the supplementary material. We ran a mixed ANOVA over percentage of correct syllables with Pause Length (Conditions 1 to 6) as within-subjects factor, and Group as between-subjects factor, including the French theta group reported in Experiment 1 (delta, theta, gamma, high gamma). The ANOVA yielded significant main effects of Group (*F*(3,92) = 42.2, *p* < .001, *η_p_²* = .579), Pause Length (*F*(4.41,406) = 42, *p* < .001, *η_p_²* = .313), and a significant interaction between the two factors (*F*(13.23,406) = 9.5, *p* < .001, *η_p_²* = .237). Given the main effect of Group, we ran one-way repeated-measures ANOVAs examining the role of Pause Length within the delta, gamma, and high gamma groups, all of which yielded a significant main effect of Pause Length (delta: *F*(5,115) = 3.6, *p* = .005, *η²* = .090; gamma: *F*(3.29,75.7) = 29.2, *p* < .001, *η²* = .347; high gamma: *F*(3.3,75.9) = 26.8, *p* < .001, *η²* = .399). The Holm corrected pairwise post hoc tests yielded the following results: No significant differences were found in the delta group (all p ≥ .093), that is, we found no evidence of accuracy changes across silence conditions. The analysis of the gamma group revealed that conditions 1–3 had significantly greater accuracy than conditions 5 and 6 (all *p* ≤ .001). Accuracy was also greater in condition 3 than in condition 4 (*p* = .006), and in condition 4 than in condition 6 (*p* = .005). In sum, accuracy decreased as silence duration augmented. A similar pattern was found in the high gamma group. Accuracy was significantly greater in the baseline than in all other conditions (all *p* ≤ .003), and in conditions 2-4 than in conditions 5-6 (all *p* ≤ .003).

**Figure 3.**
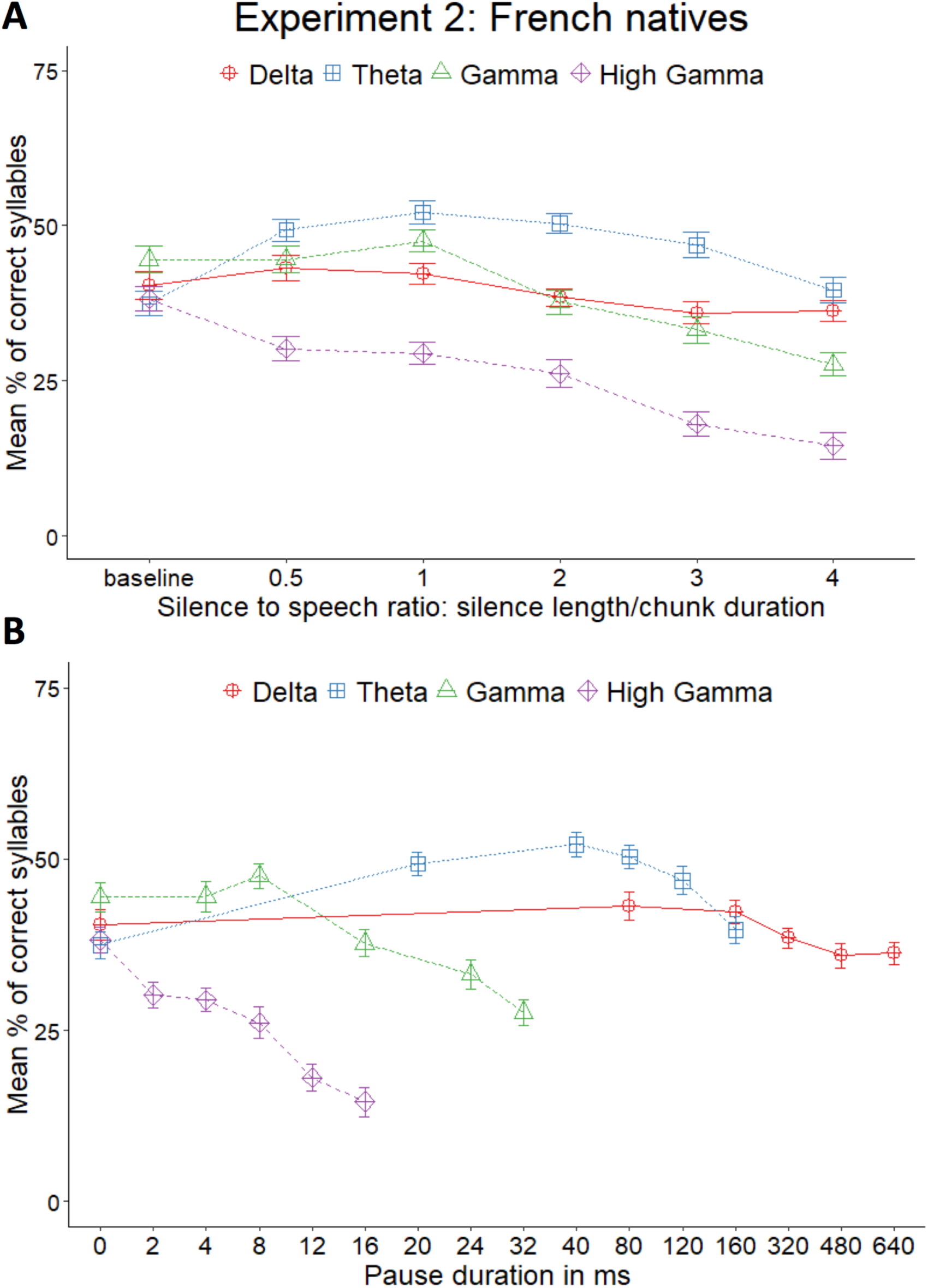
Accuracy by pause length in the four French groups. Each group’s mean accuracy is presented as a separate line. The y-axis depicts the mean percentage of correct syllables per SUS. The x-axis in panel A presents the 6 silence length conditions depicted as the proportion of silence relative to chunk duration (e.g. 0.5 entails that the pause inserted has half the duration of the speech chunk). The x-axis in panel B depicts the same results as a function of the absolute temporal durations (in ms) of the silence conditions. The Theta group is the same as French Theta in Figure 2.

## Discussion

Accelerated speech compressed to a degree that English natives no longer comprehend becomes intelligible again if silences are periodically inserted to restore the theta rhythm, a rhythm that matches the syllabic rate in natural speech and is believed to be crucial for speech comprehension^17^. Since languages show well-documented rhythmic differences^18–20^, which have observable signatures in the temporal modulations of the acoustic signal^7^, the question arises whether the rhythmic restoration effect also holds for languages that differ from English, a stress-timed language, in their linguistic rhythm. In order to test the generalizability of the English results, in Experiment 1 we tested native speakers of French and Japanese, languages that have syllable- and mora-timed rhythm, respectively.

Similarly to the English speakers in Ghitza and Greenberg^17^, we found that the periodic insertion of silences of 20, 40, 80, and 120 ms significantly improved French listeners’ comprehension as compared with no silences and silences of longer duration. Their comprehension thus showed an inverted U-shaped pattern. The fact that similar improvements in intelligibility were found in four of the six conditions evidences that the effect did not result simply from restoring the original duration of the sentences. These results suggest that, as predicted, it is restoring the theta rhythm that improves the intelligibility of accelerated speech in syllable-timed languages. While restorative effects were found in some conditions that exceed the range traditionally attributed to the theta rhythm (e.g. condition 2: 20 ms silences), this finding is unsurprising in light of the vast work probing human perception of accelerated speech, which has shown that humans, from birth, quickly adapt to accelerated speech^31^, performing well in various linguistic tasks after brief habituation to speech accelerated to 40% its original duration^29,30,32–36^. Crucially, these studies show that adults not only adapt to moderately compressed speech in their native language, but also benefit from training in unfamiliar languages as long as they belong to the same rhythmic class as the native language^30,33,35^.

We did not find such a restorative effect in the Japanese group. Rather, we found similar accuracy levels in the baseline and the 20, 40, 80 ms conditions, and a decrease for the two conditions with the longest silences (120 and 160 ms). A similar pattern was found when accuracy was measured in morae instead of syllables. Consequently, Japanese listeners did not exhibit a U-shaped pattern, but a plateau followed by a decrease in comprehension, with best performance achieved at faster rhythms as compared with French and English listeners. These results thus fit with our prediction that the highest accuracy would occur at the silence durations that best restore a given language’s basic rhythm: i.e. the syllable in French, and the mora in Japanese, in accordance with the Rhythmic Segmentation Hypothesis^11^.

Accuracy levels in the baseline condition and overall, averaging across all conditions were significantly higher in Japanese than in French (baseline: 60.5% in Japanese vs. 37.4% in French). The higher accuracy levels found in the baseline in Japanese likely result from the fact that Japanese has simpler syllabic structure and a more reduced vocalic inventory than French. French allows complex onsets and codas, while Japanese does not allow either of them, and no consonant except /n/ can occur in coda position. Also, while French has 16 vowels, Japanese only has 5, which can be short or long. As a result, Japanese has a much lower number of possible syllables than French. This smaller syllable pool increases the chance level, i.e. the probability of simply guessing a syllable correctly by chance, possibly increasing participants’ accuracy.

Ghitza and Greenberg’s^17^ study targeted the theta rhythm only. We additionally investigated whether and if yes, how the restoration of the other oscillatory rhythms impacted comprehension of French natives. In Experiment 2, we tested the delta rhythm (1–3 Hz), associated to larger prosodic phrases, and the gamma rhythm (>30 Hz), which tracks (sub)phonemic units, dividing the latter into gamma (30–80 Hz) and high gamma (>80 Hz)^1,3,28^. Unlike the theta rhythm, there was no indication of restorative effects in any of these speech rhythms. Instead, we observed a plateau across silence conditions in the delta rhythm group, and a rather linear decrease in intelligibility in both gamma conditions as silence duration increased. These results confirm our predictions, providing further evidence for the special status of the theta rhythm in speech comprehension.

In sum, we have shown that the theta rhythm is crucial for improving the comprehension of accelerated speech in a syllable-timed language, French, but not in the mora-timed language Japanese, where gains in intelligibility are instead greatest in faster rhythms. These findings thus converge with the increasing body of empirical evidence suggesting that modulation at the rate of the rhythmic unit plays a key role in understanding speech^1,7,8,25,37^. Moreover, our results support the claim that the “theta-syllable” unit is key to speech processing^37^.

## Methods

### Participants

A total of 120 participants took part in the experiment. Of these, 96 were native French speakers, while the remaining 24 participants were native Japanese speakers. The Japanese group (4 females, 20 males; mean age: 21.88 years, range 20–24 years) and 24 of the French natives (16 females, 8 males, mean age 23.25 years, range 19–32 years) formed Experiment 1’s groups. The remaining 72 French natives were randomly assigned to Experiment 2’s delta, gamma and high gamma groups (delta group: 16 females, 8 males, mean age 22.75 years, range 17–31 years; gamma group: 18 females, 6 males, mean age 24.5 years, range 19–35 years; high gamma group: 12 females, 12 males, mean age: 25.54 years, range 18–35 years). All participants were native speakers of French or Japanese, reported normal hearing and did not have language pathologies. The study was approved by the CERES ethics board (Université Paris Cité, France), and the experiments were conducted at the Integrative Neuroscience and Cognition Centre, CNRS & Université Paris Cité, Paris, France and at the University of Aizu’s Department of Computer Science and Engineering, in accordance with relevant guidelines and regulations. Informed consent was obtained from all participants, who received monetary compensation for their participation.

### Materials

Following Ghitza and Greenberg^17^, stimuli consisted of two sets—one of 96 SUS in French, one os 88 SUS in Japanese—of Semantically Unpredictable Sentences (SUS), that is, grammatically correct sentences without meaning (e.g. French: *La salle marque l’or qui vole.* “The room marks the gold that flies.”; Japanese: *Sonna e ga hen ni kogu*. “That picture rows strangely.”). The French SUS ^38^ were extracted from the corpus created by Raake and Katz^39^ (available at https://groupeaa.limsi.fr/groupeaa.limsi.fr:downloads:sus), while the Japanese SUS were created by the authors for the present experiment. All sentences were 5 to 9 syllables long and matched in syllable length between French and Japanese. They were read aloud in both languages by male native speakers, and recorded at 44.1 KHz.

Stimulus preparation followed Ghitza and Greenberg^17^ with minor deviations noted below. The recorded SUS were time-compressed by a factor of 3 using the Change Tempo function of the program Audacity®, which compresses audio files without modifying their fundamental frequency, thus avoiding compression-induced pitch shift. We then cut every compressed SUS into chunks of constant length, which varied as a function of the group (see below), and inserted silences after each chunk using Matlab®. Duration of the silences was set in advance in order to yield five periodic conditions per group (in addition to a baseline condition with no silences added). Contrary to Ghitza and Greenberg^17^, we did not include a second set of conditions with aperiodic silences, as Ghitza and Greenberg’s^17^ results indicated better performance in periodic silence rates.

Chunk and silence duration were determined on the basis of Ghitza and Greenberg^17^. In their study, they investigated the impact of restoring the theta rhythm by segmenting the SUS into 40 ms chunks and adding pauses of 20, 40, 80, 120 or 160 ms (i.e., 40 ms x 0.5, 1, 2, 3, and 4, respectively), in addition to a baseline condition with no silences added (0 ms). We used the same chunk and pause values in our French and Japanese theta groups, and set chunk duration in delta to 160 ms, gamma to 8 ms, and high gamma to 4 ms according to the following rationale. For delta, 160 ms x 3 (i.e., the duration of the uncompressed chunk) = 480 ms, which equals 2.08 Hz, falling within the delta range (1–3 Hz). Similarly, 8 ms x 3 = 24 ms, which equals 41.67 Hz, falling within the gamma range (30–80 Hz). Finally, 4 ms x 3 = 12 ms, which equals 83.33 Hz, falling within the high gamma range (>80 Hz). Silence duration in these three groups was then calculated by maintaining the same ratios relative to chunk duration as in the theta group (see Table 1). Finally, we normalized the intensity of all the SUS to 70 dB.

### Procedure

Participants were seated in front of a computer and listened to the stimuli through headphones at a volume of approximately 65 dB. The experiment was presented using the Psyscope software^40^. Prior to the experimental task, participants underwent two short trainings. Firstly, to get familiar with the task, participants listened to 8 uncompressed SUS in random order, followed by 6 compressed SUS, one per each silence condition, also presented in random order. Participants were instructed to type whatever they had understood after listening to each SUS. These SUS were not presented during the main task, contrary to Ghitza and Greenberg^17^, who trained participants with uncompressed versions of the same 96 SUS that were presented in the test. We chose not to do this in order to avoid potential memorization effects. After this short task, participants were presented with 100 utterances 3x compressed. These utterances were not SUS, but semantically correct utterances produced by 10 female French talkers or 10 male Japanese talkers, depending on the group. The goal of this second training was to allow for adaptation to compressed speech^33,41^.

During the main task, participants listened to 48 SUS divided into the 6 silence conditions, with 8 sentences per condition as in Ghitza & Greenberg^17^. Sentences from the same condition were separated by at least 4 sentences from other conditions. The 48 SUS were quasi-randomly extracted from the SUS pool so that all sentences occurred with equal frequency across participants. In addition, each sentence was randomly distributed between its conditions. That is, a single participant could only hear each SUS in a single condition. The experiment lasted about 30 minutes.

### Data analysis

Both experiments were analysed by first conducting a mix ANOVA with Group (Experiment 1: French theta, Japanese theta; Experiment 2: French delta, theta, gamma, high gamma) as a between-subjects factor, Pause Length (conditions 1 to 6) as a within-subjects factor, and percentage of correct syllables as the dependent variable. The number of syllables of the stimuli used in the study were determined on the basis of phonetic criteria, not based on their orthography. To examine the role of pause length, we subsequently run one-way repeated-measure ANOVAs for each group in both experiments, followed by Holm corrected pairwise post hoc tests for each of the groups. In Japanese, we conducted this analysis using also percentage of correct morae as the dependent variable. Following Nespor and colleagues^42^, morae were defined as sub-syllabic units that included either onset and nucleus, or a coda. Thus, syllables containing and onset and/or nucleus consisting of a single short vowel were computed as having one mora, while syllables containing an onset and/or nucleus with a long vowel or diphthong or that had a coda were considered to have two morae^43,44^.

## Acknowledgements

We wish to thank Claire Pleche and Lucie Martin for their help collecting a subset of the data, Aurélien Brest for recording the stimuli, and Prof. Reiko Mazuka for her help with coding the output of the Japanese group. This research was supported by the ECOSSud action nr. C20S02, the ERC Consolidator Grant 773202 ‘‘BabyRhythm’’, the ANR’s French Investissements d’Avenir – Labex EFL Program under Grant [ANR-10-LABX-0083], the Italian Ministry for Universities and Research FARE Grant R204MPRHKE, the European Union Next Generation EU NRRP M6C2 – Investment 2.1 - SYNPHONIA Project, as well as the Italian Ministry for Universities and Research PRIN Grant 2022WX3FM5 to Judit Gervain, Grant RYC2021-03395-I funded by MICIU/AEI/10.13039/501100011033 and, “European Union NextGenerationEU/ PRTR”, the Basque Foundation for Science Ikerbasque, and the Basque Government [IT1439-22] to Irene de la Cruz-Pavía, and the JSPS KAKENHI Grant 24K03872 to Julián Villegas.

## Author contributions

RG, JG and IdlCP conceived the experiment, JG, IdlCP and JV designed it, IdlCP, JV and CN performed the experiment, CN, KI, IdlCP and JG analyzed the data, IdlCP and JG wrote the article and all authors reviewed it.

## Data availability statement

The datasets generated during and/or analysed during the current study are available from the corresponding author Irene de la Cruz-Pavía on reasonable request.

## References

1. Ghitza, O. Linking Speech Perception and Neurophysiology: Speech Decoding Guided by Cascaded Oscillators Locked to the Input Rhythm. Front. Psychol. 2, (2011).

2. Ghitza, O., Giraud, A.-L. & Poeppel, D. Neuronal oscillations and speech perception: critical-band temporal envelopes are the essence. Front. Hum. Neurosci. 6, (2013).

3. Giraud, A.-L. & Poeppel, D. Cortical oscillations and speech processing: emerging computational principles and operations. Nat. Neurosci. 15, 511 (2012).

4. Poeppel, D. The analysis of speech in different temporal integration windows: cerebral lateralization as [] asymmetric sampling in time’. Speech Commun. 41, 245–255 (2003).

5. Poeppel, D. The neuroanatomic and neurophysiological infrastructure for speech and language. Curr. Opin. Neurobiol. 28, 142–149 (2014).

6. Schroeder, C. E., Wilson, D. A., Radman, T., Scharfman, H. & Lakatos, P. Dynamics of Active Sensing and perceptual selection. Curr. Opin. Neurobiol. 20, 172–176 (2010).

7. Varnet, L., Ortiz-Barajas, M. C., Erra, R. G., Gervain, J. & Lorenzi, C. A cross-linguistic study of speech modulation spectra. J. Acoust. Soc. Am. 142, 1976–1989 (2017).

8. Ding, N. et al. Temporal modulations in speech and music. Neurosci. Biobehav. Rev. 81, 181–187 (2017).

9. Shannon, R. V., Zeng, F.-G., Kamath, V., Wygonski, J. & Ekelid, M. Speech recognition with primarily temporal cues. Science 270, 303–304 (1995).

10. Cabrera, L. & Gervain, J. Speech perception at birth: The brain encodes fast and slow temporal information. Sci. Adv. 6, eaba7830 (2020).

11. Cutler, A. Segmentation Problems, Rhythmic Solutions. Lingua 92, 81–104 (1994).

12. Abrams, D. A., Nicol, T., Zecker, S. & Kraus, N. Right-Hemisphere Auditory Cortex Is Dominant for Coding Syllable Patterns in Speech. J. Neurosci. 28, 3958– 3965 (2008).

13. Kubanek, J., Brunner, P., Gunduz, A., Poeppel, D. & Schalk, G. The Tracking of Speech Envelope in the Human Cortex. PLOS ONE 8, e53398 (2013).

14. Ortiz Barajas, M. C., Guevara, R. & Gervain, J. The origins and development of speech envelope tracking during the first months of life. Dev. Cogn. Neurosci. 48, 100915 (2021).

15. Pefkou, M., Arnal, L. H., Fontolan, L. & Giraud, A.-L. θ-band and β-band neural activity reflects independent syllable tracking and comprehension of time-compressed speech. J. Neurosci. 37, 7930–7938 (2017).

16. Doelling, K. B., Arnal, L. H., Ghitza, O. & Poeppel, D. Acoustic landmarks drive delta–theta oscillations to enable speech comprehension by facilitating perceptual parsing. NeuroImage 85, 761–768 (2014).

17. Ghitza, O. & Greenberg, S. On the Possible Role of Brain Rhythms in Speech Perception: Intelligibility of Time-Compressed Speech with Periodic and Aperiodic Insertions of Silence. Phonetica 66, 113–126 (2009).

18. Dellwo, V. Rhythm and speech rate: A variation coefficient forΔ C. Lang. Lang.-Process. 231–241 (2006).

19. Grabe, E. & Low, E. L. Durational variability in speech and the rhythm class hypothesis. in Papers in laboratory phonology. vol. 7 515–546 (Mouton de Gruyter, Berlin, 2002).

20. Ramus, F., Nespor, M. & Mehler, J. Correlates of linguistic rhythm in the speech signal. Cognition 73, 265–292 (1999).

21. Loukina, A., Kochanski, G., Rosner, B., Keane, E. & Shih, C. Rhythm measures and dimensions of durational variation in speech. J. Acoust. Soc. Am. 129, 3258– 3270 (2011).

22. Wiget, L. et al. How stable are acoustic metrics of contrastive speech rhythm? J. Acoust. Soc. Am. 127, 1559–1569 (2010).

23. Bijeljac-Babic, R., Bertoncini, J. & Mehler, J. How do 4-day-old infants categorize multisyllabic utterances? Dev. Psychol. 29, 711–721 (1993).

24. Bertoncini, J., Floccia, C., Nazzi, T. & Mehler, J. Morae and syllables: Rhythmical basis of speech representations in neonates. Lang. Speech 38, 311–329 (1995).

25. Mehler, J., Dommergues, J., Frauenfelder, U. H. & Segui., J. The syllable’s role in speech segmentation. Cognit. Psychol. 18, 1–86 (1981).

26. Cutler, A., Mehler, J., Norris, D. & Segui, J. The Syllable’s Differing Role in the Segmentation of French and English. J. Mem. Lang. 25, 385–400 (1986).

27. Otake, T., Hatano, G., Cutler, A. & Mehler, J. Mora or Syllable? Speech Segmentation in Japanese. J. Mem. Lang. 32, 258–278 (1993).

28. Rosen, S. Temporal information in speech: acoustic, auditory and linguistic aspects. Philos. Trans. R. Soc. Lond. B. Biol. Sci. 336, 367–373 (1992).

29. Dupoux, E. & Green, K. P. Perceptual adjustment to highly compressed speech: effects of talker and rate changes. J. Exp. Psychol. Hum. Percept. Perform. 23, 914– 927 (1997).

30. Pallier, C., Sebastian-Galles, N., Dupoux, E., Christophe, A. & Mehler, J. Perceptual adjustment to time-compressed speech: A cross-linguistic study. Mem. Cognit. 26, 844–851 (1998).

31. Issard, C. & Gervain, J. Adult-like processing of time-compressed speech by newborns: A NIRS study. Dev. Cogn. Neurosci. 25, 176–184 (2017).

32. Ahissar, E. et al. Speech comprehension is correlated with temporal response patterns recorded from auditory cortex. Proc. Natl. Acad. Sci. 98, 13367–13372 (2001).

33. Mehler, J. et al. Understanding Compressed Sentences: The Role of Rhythm and Meaning a. Ann. N. Y. Acad. Sci. 682, 272–282 (1993).

34. Orchik, D. J. & Oelschlaeger, M. L. Time-compressed speech discrimination in children and its relationship to articulation. Ear Hear. 3, 37–41 (1977).

35. Sebastián-Gallés, N., Dupoux, E., Costa, A. & Mehler, J. Adaptation to time-compressed speech: Phonological determinants. Percept. Psychophys. 62, 834–842 (2000).

36. Peelle, J. E., McMillan, C., Moore, P., Grossman, M. & Wingfield, A. Dissociable patterns of brain activity during comprehension of rapid and syntactically complex speech: Evidence from fMRI. Brain Lang. 91, 315–325 (2004).

37. Aubanel, V. & Schwartz, J.-L. The role of isochrony in speech perception in noise. Sci. Rep. 10, 19580 (2020).

38. Brest, A. L’influence du rythme de la parole sur l’intelligibilité des phrases compressées : le rôle du rythme thêta. Mém. Rech. Master 2 (2019).

39. Raake, A. & Katz, B. F. SUS-based Method for Speech Reception Threshold Measurement in French.

40. Cohen, J., MacWhinney, B., Flatt, M. & Provost, J. PsyScope: An interactive graphic system for designing and controlling experiments in the psychology laboratory using Macintosh computers. Behav. Res. Methods Instrum. Comput. 25, 257–271 (1993).

41. Foulke, E. & Sticht, T. G. Review of research on the intelligibility and comprehension of accelerated speech. Psychol. Bull. 72, 50–62 (1969).

42. Nespor, M., Shukla, M. & Mehler, J. 49 Stress-timed vs. Syllable-timed Languages.

43. Liu, S. & Takeda, K. Mora-timed, stress-timed, and syllable-timed rhythm classes: Clues in English speech production by bilingual speakers. (2021) doi:10.1556/2062.2021.00469.

44. Kubozono, H. I Introduction to Japanese phonetics and phonology. in Handbook of Japanese Phonetics and Phonology (ed. Kubozono, H.) 1–40 (De Gruyter Mouton, 2015). doi:10.1515/9781614511984.1.

