## Supplementary information for "The Role of Syllabic Rhythm in Speech Perception Across Languages"

**Table S1.** Mean percentage and standard error of correct syllables in French and Japanese, as well as morae in Japanese, divided by group and condition. The French theta and Japanese theta groups were tested in Experiment 1, while the French delta, gamma and high gamma groups were tested in Experiment 2.

| | French | | | | Japanese | |
|---|---|---|---|---|---|---|
| | **Delta** | **Theta** | **Gamma** | **High gamma** | **Theta (syllables)** | **Theta (morae)** |
| **Condition 1: baseline** | 40.4 ±2.24 | 37.4 ±2.00 | 44.5 ±2.15 | 38.2 ±1.92 | 60.5 ±1.75 | 64.2 ±1.75 |
| **Condition 2** | 43.1 ±2.06 | 49.3 ±1.77 | 44.6 ±2.19 | 30.2 ±1.96 | 62.6 ±2.69 | 65.6 ±2.62 |
| **Condition 3** | 42.3 ±1.73 | 52.1 ±1.85 | 47.5 ±1.83 | 29.4 ±1.74 | 55.1 ±2.60 | 59.5 ±2.32 |
| **Condition 4** | 38.5 ±1.40 | 50.3 ±1.65 | 37.7 ±1.94 | 26.1 ±2.27 | 53.3 ±2.92 | 57.6 ±2.79 |
| **Condition 5** | 35.9 ±1.77 | 46.8 ±2.06 | 33.2 ±2.11 | 18.1 ±1.99 | 48.1 ±3.18 | 52.3 ±3.16 |
| **Condition 6** | 36.3 ±1.65 | 39.6 ±2.01 | 27.6 ±1.85 | 14.6 ±2.16 | 42.4 ±2.66 | 48.4 ±2.69 |

**Table S2.** Experiment 1 and 2’s post-hoc analyses consisting of Holm-corrected pairwise comparisons. Only significant comparisons are reported. The upper table compares the accuracy of Experiment 1’s French vs. Japanese theta groups, in each of the silence durations (presented in ms). The lower table depicts the results of the French delta, theta, gamma and high gamma groups (presented as ratio silence/speech).

<table>
<tr><th colspan="3">Experiment 1:<br>theta rhythm in French vs. Japanese</th></tr>
<tr><th>Pause length</th><th colspan="2">p-value</th></tr>
<tr><td>Baseline (condition 1)</td><td colspan="2">< .001</td></tr>
<tr><td>20 ms (condition 2)</td><td colspan="2">< .001</td></tr>
<tr><td colspan="3"></td></tr>
<tr><th colspan="3">Experiment 2:<br>delta, theta, gamma, and high gamma rhythms in French</th></tr>
<tr><th>Ratio<br>silence/speech</th><th>Groups</th><th>p-value</th></tr>
<tr><td rowspan="3">0.5<br>(condition 2)</td><td>delta vs. high gamma</td><td>< .001</td></tr>
<tr><td>theta vs. high gamma</td><td>< .001</td></tr>
<tr><td>gamma vs. high gamma</td><td>< .001</td></tr>
<tr><td rowspan="4">1<br>(condition 3)</td><td>delta vs. theta</td><td>= .001</td></tr>
<tr><td>delta vs. high gamma</td><td>< .001</td></tr>
<tr><td>theta vs. high gamma</td><td>< .001</td></tr>
<tr><td>gamma vs. high gamma</td><td>< .001</td></tr>
<tr><td rowspan="5">2<br>(condition 4)</td><td>delta vs. theta</td><td>< .001</td></tr>
<tr><td>delta vs. high gamma</td><td>< .001</td></tr>
<tr><td>theta vs. gamma</td><td>< .001</td></tr>
<tr><td>theta vs. high gamma</td><td>< .001</td></tr>
<tr><td>gamma vs. high gamma</td><td>< .001</td></tr>
<tr><td rowspan="3">3<br>(condition 5)</td><td>delta vs. theta</td><td>= .001</td></tr>
<tr><td>delta vs. high gamma</td><td>< .001</td></tr>
<tr><td>theta vs. gamma</td><td>< .001</td></tr>
</table>

| | | |
|---|---|---|
| | theta vs. high gamma | < .001 |
| | gamma vs. high gamma | < .001 |
| 4 (condition 6) | delta vs. gamma | = .012 |
| | delta vs. high gamma | < .001 |
| | theta vs. gamma | < .001 |
| | theta vs. high gamma | < .001 |
| | gamma vs. high gamma | < .001 |
